# USP10 Facilitates Homologous Recombination-Mediated DNA Double-Strand Break Repair through Localization to the Nucleolus

**DOI:** 10.64898/2026.05.07.723676

**Authors:** Koichi Utani, Ryo Sakasai, Toshiki Himeda, Takako Okuwa, Kuniyoshi Iwabuchi, Masaya Higuchi

**Affiliations:** Department of Microbiology Kanazawa Medical University, Uchinada, Kahoku, Ishikawa, Japan; Department of Biochemistry I, Kanazawa Medical University, Uchinada, Kahoku, Ishikawa, Japan

**Keywords:** DNA Breaks, Double-Stranded, DNA Repair, Nucleolus, Recombination, Genetic, Ubiquitin-Specific Proteases

## Abstract

Ubiquitin-specific protease 10 (USP10) is a multifunctional deubiquitinating enzyme that primarily regulates cellular stress responses, including the DNA damage response. Here, we show that USP10 is required for homologous recombination (HR)-mediated repair of DNA double-strand breaks (DSBs) and for the maintenance of genomic stability. USP10-depleted cells exhibit spontaneous micronuclei, impaired DSB repair following zeocin and camptothecin treatment, and reduced sister chromatid exchange. These cells are also more sensitive to irradiation and mitomycin C and display increased chromosomal abnormalities after mitomycin C treatment. Persistent RAD51 foci formation in USP10-depleted cells suggests that USP10 functions at a step downstream of RAD51 nucleofilament formation. This function of USP10 in facilitating HR repair depends on deubiquitinase activity but is independent of G3BP1/2 and PABP binding. In addition, a newly identified nucleolar localization signal is required for USP10’s function in DSB repair. Together, these findings indicate that USP10 maintains genome integrity by localizing to the nucleolus and facilitating HR-mediated repair of DSBs.

## 1 Introduction

Ubiquitination and deubiquitination of intracellular proteins are essential post-translational modifications that regulate protein abundance, quality, and function. Dysregulation of the ubiquitination/deubiquitination system has been implicated in the pathogenesis of various diseases, including cancer and neurodegenerative disorders (1). Ubiquitin-specific protease 10 (USP10) is a multifunctional deubiquitinating enzyme with diverse biological roles (2). Under various stress conditions, USP10 is recruited to stress granules (SGs) together with the SG-nucleating factor G3BP1/2, where it regulates SG assembly (3). In addition, USP10 has been reported to modulate autophagy, energy metabolism, and protein translation in response to stress (4-7). Beyond these functions, USP10 contributes to various DNA repair and damage response processes, including translesion DNA synthesis (8), mismatch repair (9), and p53 regulation (10). Collectively, these findings suggest that USP10 has evolved as a stress-responsive deubiquitinating enzyme that maintains cellular homeostasis under various stress conditions, including DNA damage.

Among the different types of DNA damage, DNA double-strand breaks (DSBs) are the most deleterious lesions because they can result in nucleotide deletions or insertions and, if left unrepaired, lead to chromosomal rearrangements such as translocations—events that can promote tumorigenesis (11). Mammalian cells repair DSBs primarily through two pathways: non-homologous end joining (NHEJ) and homologous recombination (HR) (12). In NHEJ, DSB ends are directly ligated after limited end processing, which can introduce small insertions or deletions, rendering it an error-prone repair pathway (13). In contrast, HR utilizes an undamaged sister chromatid as a repair template, ensuring accurate, error-free DSB repair (14).

During HR, end resection at the DSB site generates a ssDNA nucleofilament that invades the homologous duplex DNA to initiate homology search, strand annealing, and DNA synthesis, ultimately forming a displacement loop (D-loop). The repair process is then completed through synthesis-dependent strand annealing (SDSA), Holliday junction (HJ) resolution, or HJ dissolution (14). Successful execution of HR requires multiple post-translational modifications, including ubiquitination and deubiquitination of HR-related proteins (15, 16). Several deubiquitinases have been reported to regulate HR by modulating the stability and activity of these repair factors (17-20). However, because HR is a multistep process involving numerous regulatory proteins, the overall landscape of ubiquitin signaling in HR remains incompletely understood. Identifying additional deubiquitinases and their substrates involved in HR is therefore essential for elucidating the molecular mechanisms governing this repair pathway.

In this study, we investigate the function of USP10 in the DSB repair and show that it promotes HR-mediated DSB repair. We further demonstrate that its newly identified nucleolar localization signal (NoLS) is crucial for USP10’s HR-facilitating function.

## 2 Materials and Methods

Materials and Methods used in this study include Cell culture; Plasmids; Lentiviral transduction; Establishment of USP10 knockout or knockdown cell lines; DNA damage induction; Antibodies; Immunostaining; Western blotting; Neutral comet assay; MTS cell viability assay; Clonogenic colony formation assay; Sister chromatid exchange assay; Analysis of chromosomal aberrations; Statistical analysis. Detailed information is shown in Doc. S1.

## 3 Results

### 3.1 USP10 Depletion Induces Genomic Instability

To investigate the role of USP10 in DNA damage repair, we first generated USP10-KO mouse embryonic fibroblasts (MEFs) (Figure S1) and quantified micronuclei formation—a hallmark of genomic instability (21)—compared with the level in WT MEFs. USP10-KO MEFs displayed a significantly higher frequency of micronuclei compared with WT cells, indicating that USP10 loss promotes genomic instability (Figure 1A,B). Consistent with this result, neutral comet assays revealed that USP10-KO MEFs accumulated more spontaneous DNA DSBs than WT MEFs did (Figure 1C,D). Notably, these micronuclei were often positioned adjacent to the nucleus, a feature characteristic of micronuclei derived from chromosome bridges and breaks (22). In line with this observation, USP10-KO MEFs exhibited a higher incidence of chromosome bridges in interphase cells (Figure 1E,F). Together, these findings demonstrate that USP10 is critical for preserving genomic stability, particularly by facilitating the repair of DNA DSBs.

**FIGURE 1.**
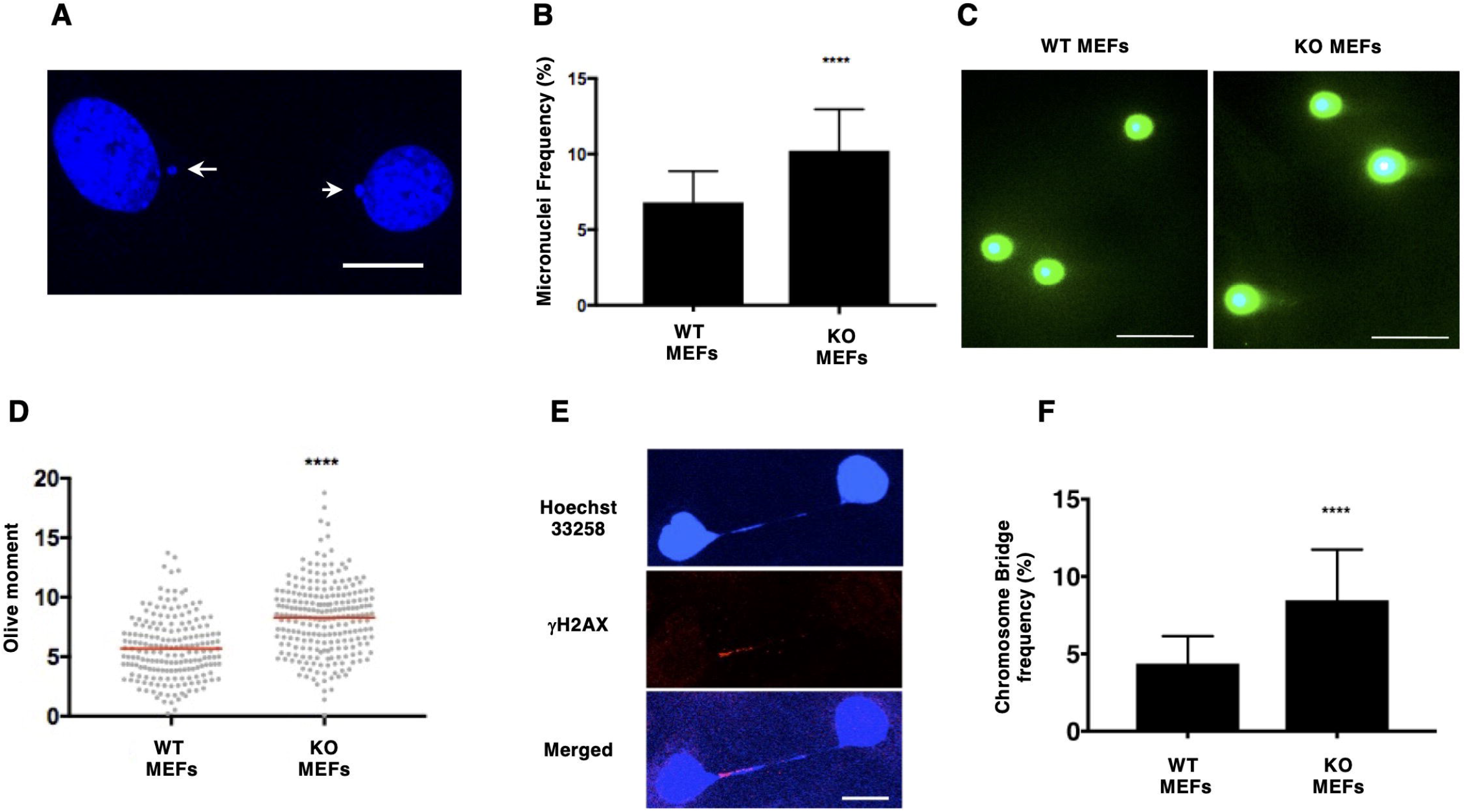
USP10-KO mouse embryonic fibroblasts (MEFs) exhibit genomic instability. (A) Micronuclei formation (arrows) in USP10-KO MEFs. Nuclei were stained with Hoechst 33258. Scale bar, 20 μm. (B) Frequency of micronuclei formation in WT and USP10-KO MEFs. At least 5,000 cells per genotype were evaluated. (C,D) Neutral comet assays performed in WT and USP10-KO MEFs. Representative images (C) and quantification of Olive tail moments in WT and USP10-KO MEFs (D). At least 120 cells per genotype were analyzed. Scale bar, 50 μm. (E) Chromosome bridges in USP10-KO MEFs stained with γH2AX antibody and Hoechst 33258. Scale bar, 20 μm. (F) Frequency of chromosome bridge in WT and USP10-KO MEFs. At least 1,800 cells per genotype were analyzed. *P* values were determined by the Mann–Whitney *U* test. ****, *P* < 0.0001.

### 3.2 USP10 Facilitates DSB repair in a Deubiquitinase-Dependent Manner

To assess whether USP10 participates in DNA DSB repair, we monitored γH2AX foci formation, a well-established marker of DNA DSBs, following treatment with zeocin, which intercalates into DNA and induces DSBs. In the absence of zeocin, USP10-KO MEFs exhibited higher numbers of γH2AX foci than WT cells, consistent with the results shown in Figure 1. Upon zeocin treatment, γH2AX foci were induced to comparable levels in both WT and USP10-KO MEFs. However, whereas γH2AX foci returned to basal levels 9 h after zeocin removal in WT MEFs, they remained elevated in USP10-KO MEFs (Figure 2A,B). Neutral comet assays further confirmed impaired DSB repair in USP10-KO MEFs (Figure 2C). Importantly, reintroduction of mouse USP10 (both full length [FL] and FL-HA tagged [FL-HA] USP10) into USP10-KO MEFs restored DSB repair capacity, as demonstrated by γH2AX staining and neutral comet assays (Figure 2D–G). USP10 is known to associate with G3BP1/2 and PABP through its N-terminal FGDF and PAM2 motifs, respectively (3). However, deletion of these interaction domains (HA-Δ95) did not affect its ability to support DSB repair. In contrast, expression of a catalytically inactive deubiquitinase mutant (HA-C418A) failed to rescue repair activity (Figure 2D–G). These findings indicate that USP10 promotes DSB repair in a manner dependent on its deubiquitinase activity but independent of its interaction with G3BP1/2 and PABP.

**FIGURE 2.**
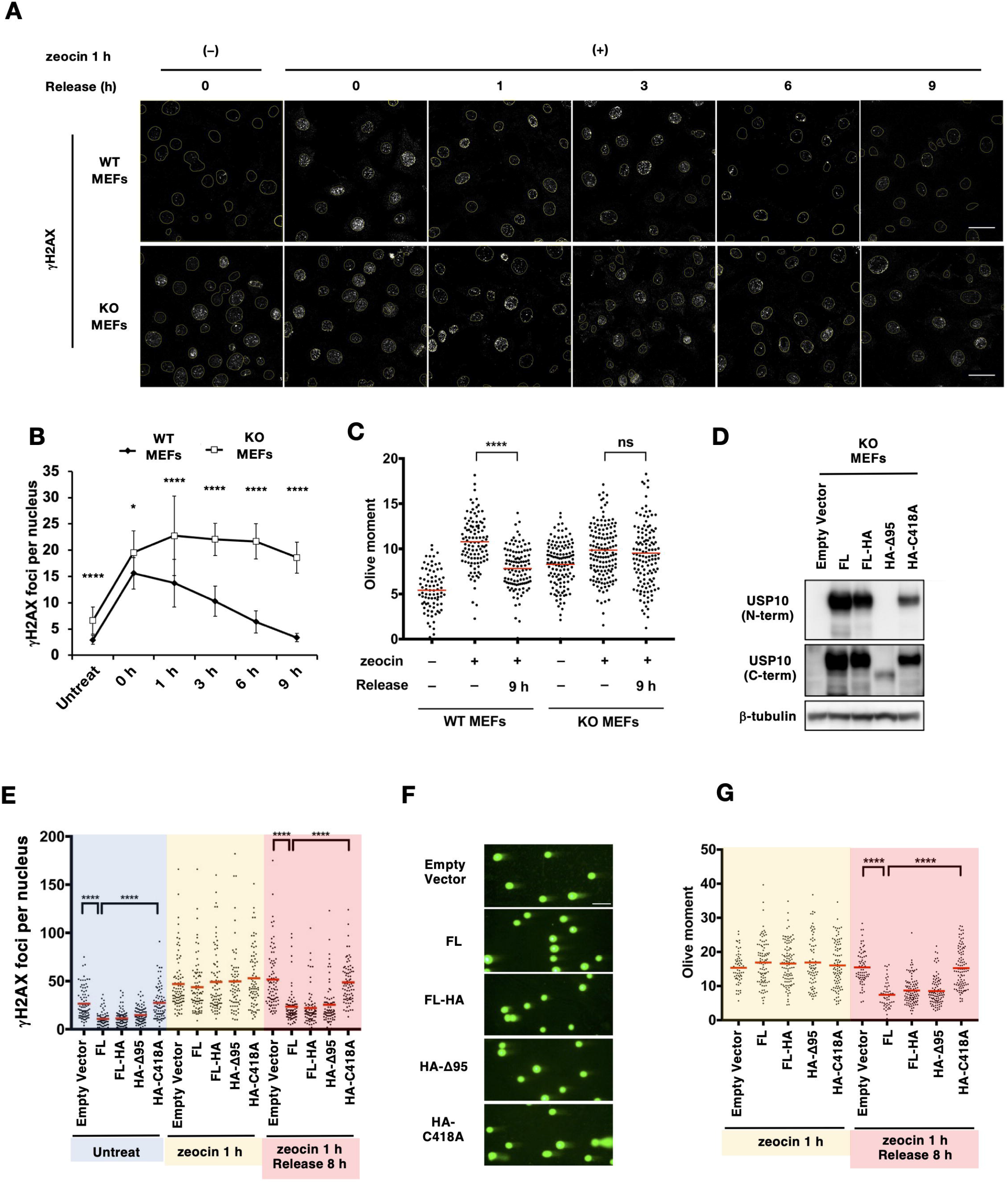
USP10 is required for double-strand break (DSB) repair in zeocin-treated MEFs. (A) Time course of γH2X foci formation before and after zeocin treatment in WT and USP10-KO MEFs. Cells were treated with 100 μg/mL zeocin for 1 h, washed, and incubated in drug-free medium for the indicated times. Nuclei were counterstained with Hoechst 33258 and are outlined by yellow lines. (B) Quantification of γH2AX foci in (A). At least 100 cells per genotype were analyzed at each time point. (C) Neutral comet assays before and after 1 h zeocin treatment followed by 9 h release. (D) Expression of reintroduced USP10 and its mutants in USP10-KO MEFs. Two antibodies recognizing either the N-terminus (N-term) or the C-terminus (C-term) of USP10 were used for western blotting (WB). β-tubulin was used as a loading control. (E) γH2X foci formation in USP10-KO MEFs expressing WT or USP10 mutants before and after 1 h zeocin treatment and 8 h release. (F,G) Neutral comet assays in USP10-KO MEFs expressing WT or USP10 mutants after 1 h zeocin treatment and 8 h release. (G) Quantification of the experiment in (F). Scale bar, 50 μm. *P* values were determined by the Mann–Whitney *U* test (B,C) and Dunnett’s multiple comparison test (E,G). ns, not significant; ****, *P* < 0.0001.

To further investigate USP10’s role in human cells, we depleted USP10 in HCT116 (colon cancer), MCF-7 (breast cancer), and WI-38 (normal fibroblast) cells. In all cases, USP10 knockdown (KD) resulted in defective DSB repair following zeocin treatment (Figure S2A–D). Reintroduction of WT human USP10, but not the catalytically inactive C424A mutant, into USP10-depleted HCT116 cells restored repair capacity, confirming the requirement for USP10’s deubiquitinase activity in human cells as well (Figure S2E,F). In addition, four independent USP10-KO clones of HCT116 generated using CRISPR-Cas9 (Figure S2G) exhibited a significant increase in micronuclei frequency (Figure S2H) and persistent γH2AX foci after zeocin treatment followed by a 7-h recovery period (Figure S2I). Collectively, these results establish USP10 as a key factor required for efficient DSB repair in both mouse and human cells, acting through its deubiquitinase activity.

### 3.3 USP10 Facilitates HR-Mediated DSB Repair Pathway

DNA DSBs are repaired primarily through two distinct pathways: the error-prone NHEJ pathway and the error-free HR pathway. To determine which pathway predominantly repairs zeocin-indued DNA damage, we first examined the effects of SCR7, an inhibitor of the essential NHEJ factor DNA ligase IV in MEF cells. SCR7 reduced cell viability and DSB repair capacity only in etoposide (ETP)-treated cells, consistent with the fact that ETP-induced DSBs are primarily resolved via NHEJ (Figure 3A,B) (23). In contrast, SCR7 had no effect on cell viability or repair efficiency in cells treated with zeocin or camptothecin (CPT). Interestingly, SCR7 even increased repair capacity in CPT-treated cells. Since CPT-induced DSBs are mainly repaired by HR (24), these findings suggest that zeocin-induced DSBs are also primarily repaired through the HR pathway. This conclusion is further supported by previous work demonstrating that zeocin-induced γH2AX foci colocalize with RAD51 foci, indicative of ssDNA nucleofilament formation—a critical step in HR repair (25). Thus, the defective repair of zeocin-induced DSBs in USP10-KO MEFs appears to result from impaired HR repair.

**FIGURE 3.**
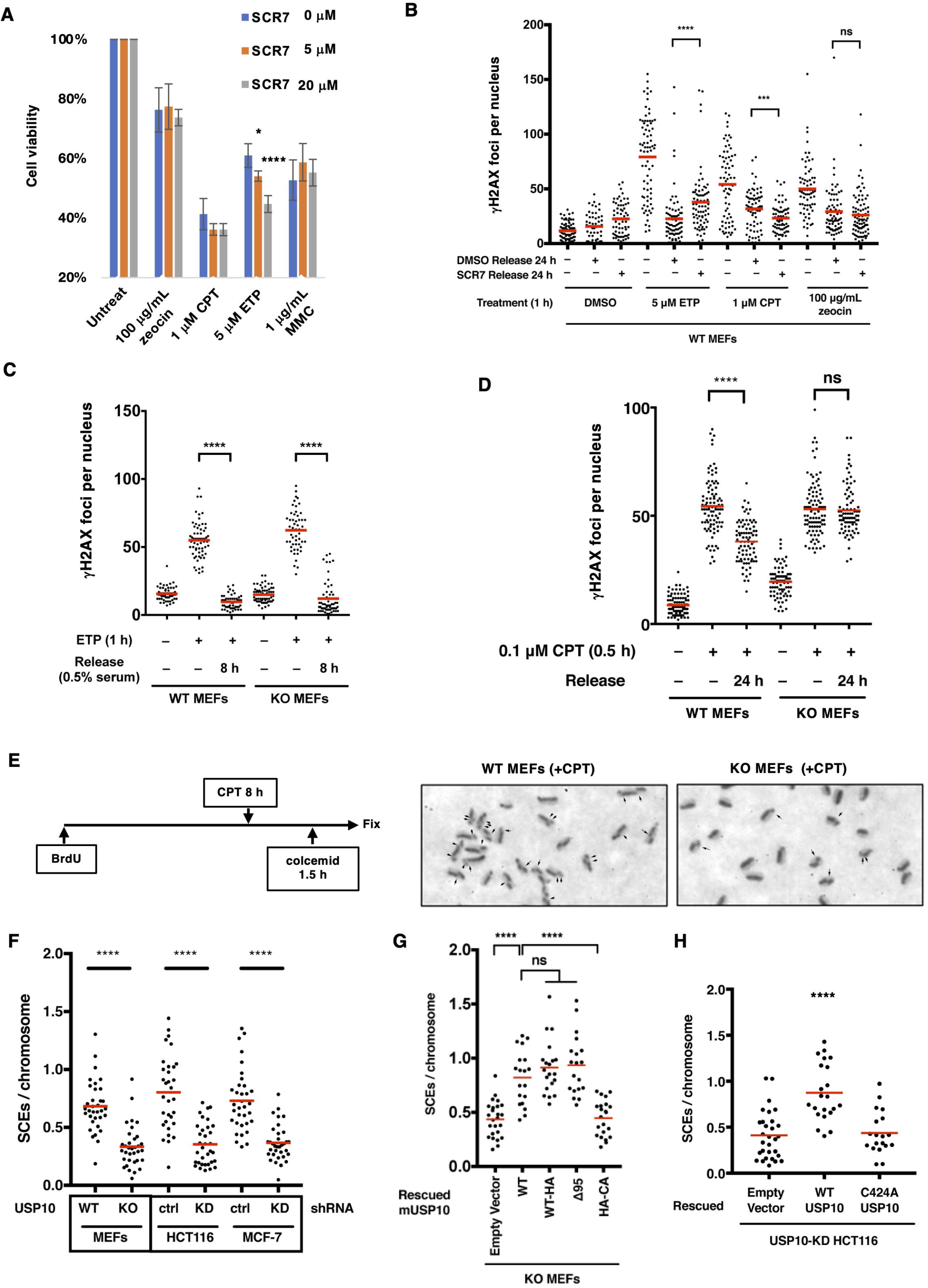
USP10 is required for DSB repair mediated by homologous recombination (HR) but not non-homologous end joining (NHEJ). (A) Effect of SCR7 on survival of MEFs treated with zeocin, camptothecin (CPT), etoposide (ETP), or mitomycin C (MMC). WT MEFs were treated with indicated drugs in the presence or absence of SCR7 for 30 h. Cell survival was measured by MTS assay. (B) Effect of SCR7 on DSB repair assessed using γH2AX foci after drug treatment and release. At least 60 cells were analyzed per condition. (C) γH2AX foci in WT and USP10-KO MEFs synchronized in G1 phase of the cell cycle. Cells were synchronized in G1 by culturing in medium containing 0.5% FCS for 24 h, treated with ETP, and released in drug-free medium containing 0.5% FCS before γH2AX foci quantification. At least 50 cells were analyzed per genotype. (D) γH2AX foci in WT and USP10-KO MEFs before and after treatment with CPT followed by 24 h release. At least 80 cells per genotype were analyzed. (E) Sister chromatid exchange (SCE) assay schematic and representative metaphase spreads. Arrowheads indicate SCE sites. SCE frequencies in WT and USP10-KO MEFs, control (ctrl) and USP10-knockdown (KD) HCT116 and MCF-7 cells (F), USP10-KO MEFs expressing WT or indicated USP10 mutants (G), and USP10-KD HCT116 cells expressing WT or indicated USP10 mutants (H). Thirty metaphase spreads were analyzed per sample. *P* values were determined by the Mann–Whitney *U* test (B–D,F) and Dunnett’s multiple comparison test (A,G,H). ns, not significant; *, *P* < 0.05; ****, *P* < 0.0001.

To confirm that USP10-KO MEFs are defective in HR but not NHEJ, we first assessed NHEJ activity by monitoring ETP-induced γH2AX foci formation in G0/G1-phase serum-starved MEFs, where NHEJ is the only available repair pathway. No significant differences in DNA repair activity were observed between WT and USP10-KO MEFs, indicating that NHEJ remains intact in the absence of USP10 (Figure 3C). By contrast, CPT-induced DSBs failed to be repaired in USP10-KO MEFs, similar to the phenotype observed with zeocin treatment, further supporting the conclusion that USP10 deficiency specifically impairs the HR pathway (Figure 3D).

To directly measure HR pathway activity, we performed a sister chromatid exchange (SCE) assay (Figure 3E). USP10-KO MEFs displayed a significantly lower SCE frequency than WT MEFs did (Figure 3F). Similarly, USP10 KD in HCT116 and MCF-7 suppressed SCE frequency. Reintroduction of WT and Δ95 USP10, but not the C418A mutant, restored the SCE frequency in USP10-KO MEFs (Figure 3G). The USP10-KD HCT116 introduced with WT-USP10 but not the C424A mutant showed the same result (Figure 3H). Together, these results demonstrate that USP10’s deubiquitinase activity is essential for efficient HR-mediated repair of DSBs.

### 3.4 USP10 Depletion Sensitizes Cells to DSB-Inducing Agents

We next examined the effect of USP10 depletion on cell viability following treatment with various DSB-inducing agents. WT and USP10-KO MEFs were treated with zeocin, ETP, CPT, or mitomycin C (MMC), and cell viability was assessed using an MTS assay (Figure 4A). USP10-KO MEFs exhibited increased sensitivity to all tested agents compared with WT cells, consistent with impaired DSB repair in the absence of USP10.

**FIGURE 4.**
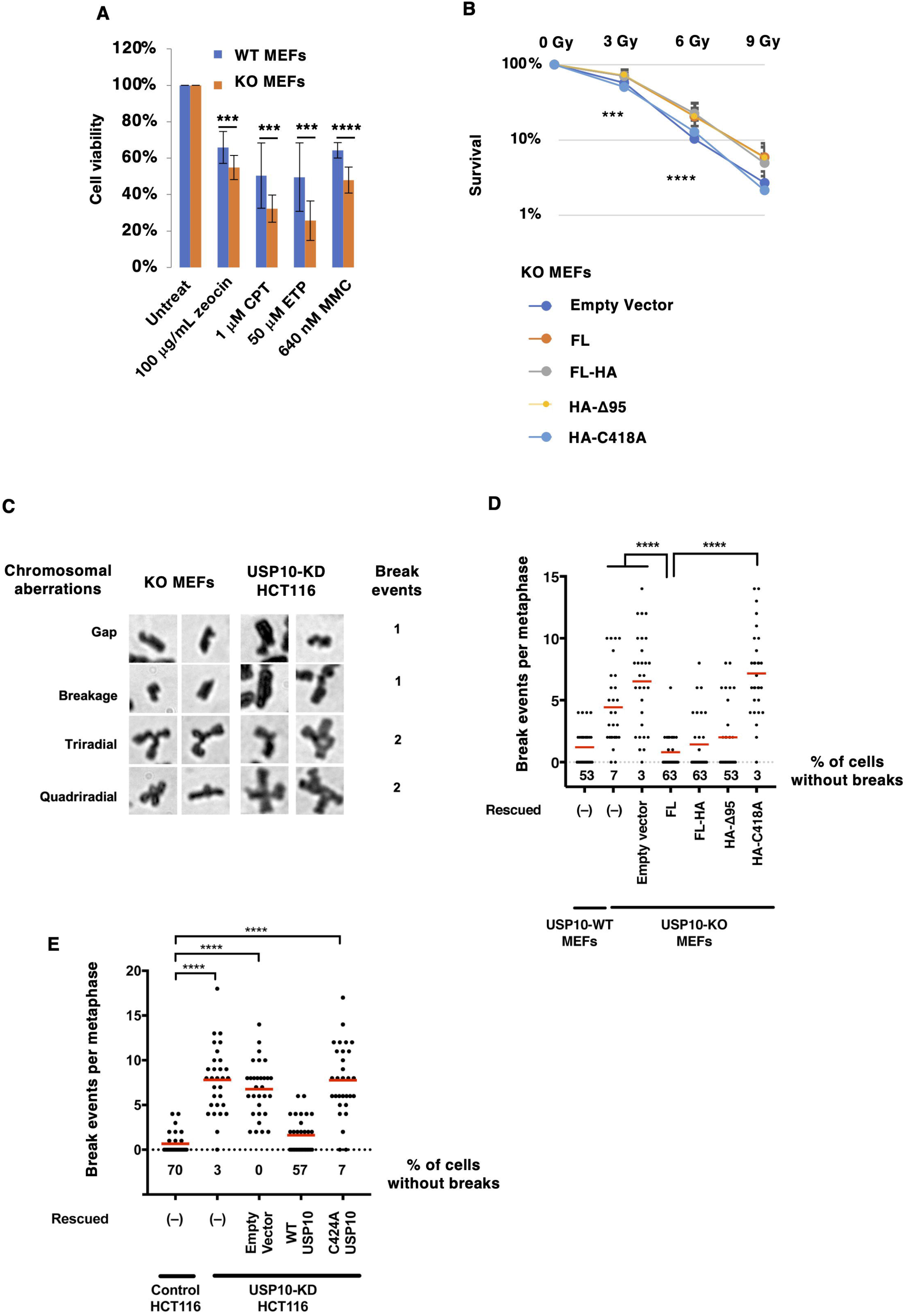
USP10-depleted cells are more sensitive to DNA damage. (A) Survival of WT and USP10-KO MEFs following treatment with DNA-damaging agents. MEF cells were treated with the indicated drugs for 30 h, and cell viability was measured by MTS assay. (B) Clonogenic colony formation assays of USP10-KO MEFs expressing WT or indicated USP10 mutants following X-ray irradiation. (C) Representative images of chromosomal aberrations in USP10-KO MEFs and USP10-KD HCT116 cells following MMC treatment. Chromatid gaps or breaks were scored as single-break events, whereas triradial and quadriradial chromosomes were scored as double-break events. (D,E) Quantification of chromosomal aberrations following MMC treatment in WT, USP10-KO, and USP10-KO MEFs expressing WT or the indicated USP10 mutants (D), and in control, USP10-KD HCT116, and USP10-KD HCT116 cells expressing WT or the indicated USP10 mutants (E). The percentages of cells without aberrations are indicated. *P* values were determined by Student’s *t* test (A,B) and Dunnett’s multiple comparison test (D,E). ***, *P* < 0.001; ****, *P* < 0.0001.

To further assess DNA damage sensitivity, we performed clonogenic survival assays following irradiation. USP10-KO MEFs exhibited a mild but significant reduction in colony formation (Figure S3A), which was rescued by the reintroduction of WT USP10 and the Δ95 mutant but not the catalytically inactive C418A mutant (Figure 4B). A similar decrease in clonogenic survival was observed in USP10-KO MEF cells following MMC treatment (Figure S3B).

The observed MMC sensitivity in USP10-depleted cells prompted us to investigate MMC-induced chromosomal abnormalities. Following an established protocol (26), we quantified chromosomal aberrations, scoring chromosome gaps and breaks as single-break events, and triradial and quadriradial structures as double-break events (Figure 4C). Both USP10-KO MEFs and USP10-KD HCT116 cells exhibited a marked increase in chromosomal abnormalities after MMC treatment. Reintroduction of FL USP10 and the Δ95 mutant, but not the catalytically inactive mutant, restored chromosomal integrity to normal levels (Figure 4D,E). This phenotype in USP10-depleted cells is likely attributable to defective HR repair, which is essential for the resolving the DSBs generated during interstrand crosslink (ICL) repair. Alternatively, USP10 depletion may directly impair the removal of ICLs.

### 3.5 Persistent RAD51 Foci Formation in USP10-KO Cells

HR repair proceeds through several sequential steps; DSB end resection generating 3’-ssDNA, RAD51 loading onto ssDNA and filament formation, strand invasion and D-loop formation, RAD51 dissociation and DNA synthesis, and finally the resolution or dissolution of joint DNA molecules (14). To determine which step of HR is impaired in USP10-KO cells, we examined the formation of RAD51 foci, as RAD51 is a central mediator of HR repair that forms nuclear foci upon loading onto resected DSB ends. In WT HCT116 cells, RAD51 accumulated as distinct foci 3 h after irradiation, and the number of foci decreased by 9 h, consistent with the completion of HR-mediated repair (Figure 5A–D). In contrast, USP10-KO cells retained high levels of RAD51 foci even 9 h after irradiation, with a subset of these foci overlapping with persistent γH2AX foci. Similar results were observed in zeocin-treated cells (Figure S4). These findings suggest that RAD51 loading onto resected ssDNA occurs normally in USP10-KO cells, but a subsequent step of HR—most likely strand invasion or RAD51 dissociation prior to DNA synthesis—is impaired.

**FIGURE 5.**
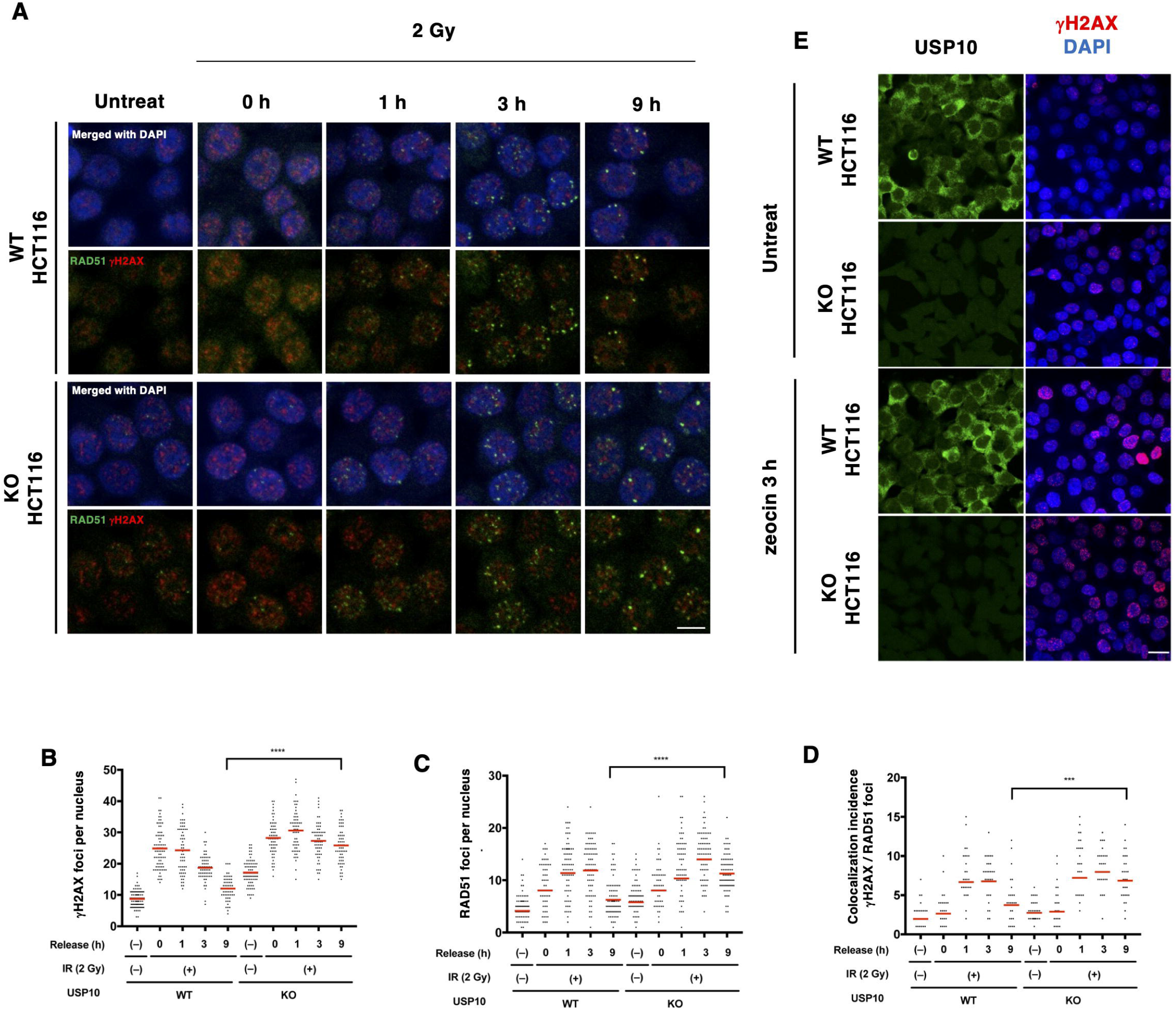
Persistent RAD51 foci formation in USP10-KO HCT116 cells. (A) RAD51 (green) and γH2AX (red) foci in WT and USP10-KO HCT116 cells at the indicated time points following X-ray irradiation. Nuclei were counterstained with DAPI. Scale bar, 10 μm. (B–D) Quantification of RAD51 (B), γH2AX (C), and colocalized RAD51/γH2AX foci (D) shown in (A). (E) Localization of USP10 (green) and γH2AX (red) in WT and USP10-KO HCT116 cells before and after zeocin treatment. Nuclei were counterstained with DAPI. Scale bar, 20 μm. *P* values were determined by the Mann–Whitney *U* test. ****, *P* < 0.0001.

To investigate whether USP10 contributes to the HR repair process by directly deubiquitinating HR-related proteins at DSB sites, we examined the localization of USP10 following DSB induction. In zeocin-treated HCT116 cells, USP10 was primarily localized in the cytoplasm before treatment and did not colocalize with γH2AX after treatment (Figure 5E). To further assess USP10 localization, we analyzed its distribution in UV-irradiated cells following BrdU labeling, which induces DSBs throughout the nucleus (27). Again, we did not observe any significant relocalization of USP10 to the nucleus (Figure S5). These findings suggest that USP10 is unlikely to directly participate in the HR repair process at DSB sites. Instead, it may facilitate HR repair by indirectly regulating steps downstream of RAD51 recruitment.

### 3.6 USP10 Region Spanning Amino Acids 139–166 Is Required for HR Repair

We next sought to identify which region of USP10, apart from its deubiquitinase domain, is essential for HR repair. As shown in Figure 2, deletion of the N-terminal 95 amino acids (Δ95 mutant) did not impair DSB repair capacity. According to the UniProt database, the N-terminal region of USP10 contains three intrinsically disordered segments spanning amino acids 139–166, 194–257, and 307–337, which may be functionally important. Given this, we first assessed the contribution of the 139–166 region, as it is particularly rich in lysine and arginine residues. To this end, we generated a double-deletion mutant lacking the first 95 amino acids and residues 139–166 (Δ95/Δ139–166 mutant) (Figure 6A). This construct was introduced into USP10-KO HCT116 cells, and DSB repair capacity was evaluated following zeocin treatment. While the Δ95 mutant efficiently repaired DSBs, the Δ95/Δ139–166 mutant failed to do so (Figure 6B). The Δ95/Δ139–166 mutant did not show an increase in SCE frequency after CPT treatment (Figure 6C) or reduced abnormal mitotic chromosome formation after MMC treatment (Figure 6D). Moreover, deletion of only the 139–166 region from FL USP10 also abrogated DSB repair capacity following zeocin treatment (Figure 6E). We verified that the Δ139–166 mutant retained normal deubiquitinase activity, as ribosomal protein S3 (RPS3) was efficiently deubiquitinated in cells expressing this mutant (Figure 6F) (7). These findings indicate that the 139–166 region of USP10 is essential for HR repair.

**FIGURE 6.**
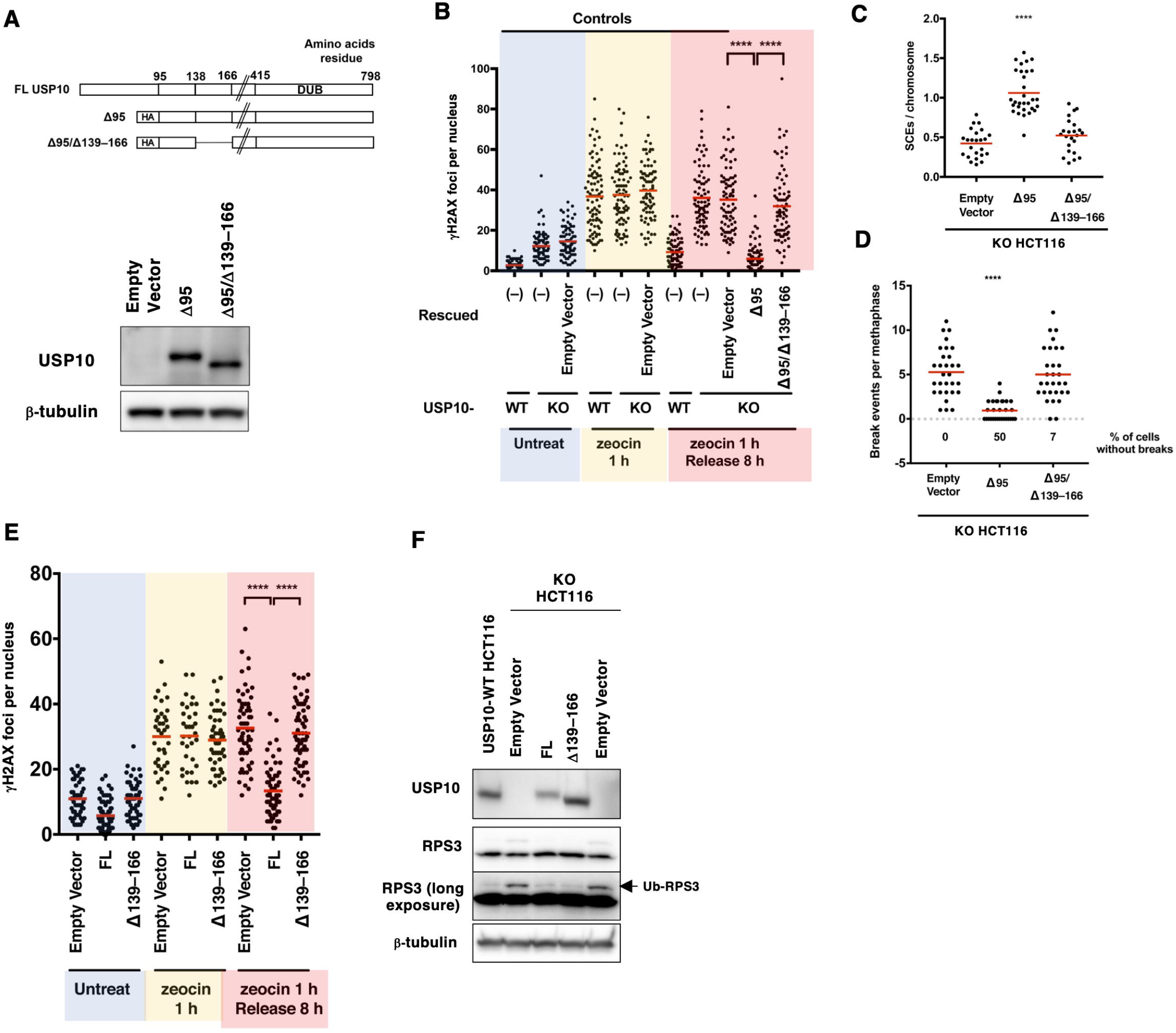
The USP10 139–166 region is required for HR repair. (A) Expression of USP10 mutants determined by WB using an anti-USP10 antibody. (B–D) DSB repair assessed using γH2AX foci following zeocin treatment (B), SCE frequencies following CPT treatment (C), and chromosomal aberration following MMC treatment (D) in USP10-KO HCT116 cells expressing Δ95 or Δ95/Δ139–166 USP10 mutants. (E) DSB repair in USP10-KO HCT 116 cells expressing FL USP10 or Δ139–166 USP10 mutant following zeocin treatment. (F) Ribosomal protein S3 (RPS3) ubiquitination in indicated cells determined by WB using anti-RPS3 and anti-USP10 antibodies. Monoubiquitinated RPS3 is indicated by an arrow. *P* values were determined by Dunnett’s multiple comparison test. ****, *P* < 0.0001.

We next examined whether regions downstream of residue 166, up to the deubiquitinase domain, are also required for DSB repair. To this end, we generated a series of deletion mutants covering this interval. For potential use in future proximity-labeling studies, we fused AirID (28), a biotinylating enzyme, to the N terminus of these constructs (Figure S6A). To confirm that AirID fusion itself does not interfere with USP10 function in DSB repair, we first tested AirID-Δ138 and AirID-Δ166 mutants. AirID-Δ138 USP10 restored DSB repair capacity, whereas AirID-Δ166 did not, consistent with the requirement of the 139–166 region (Figure S6A). We then analyzed additional deletion mutants spanning regions between residue 166 and the deubiquitinase domain. All of these mutants retained DSB repair capacity comparable to that of the Δ138 mutant, except for the Δ338–415 construct, which was highly unstable (Figure S6B). These results indicate that, apart from the deubiquitinase domain, the 139–166 region is the only segment of USP10 that is essential for HR-mediated DSB repair.

### 3.7 The USP10 139–166 Region Contains a Putative NoLS

As noted above, the 139–166 region of USP10 is particularly rich in lysine and arginine residues, a common feature of nuclear localization signals (NLSs). Consistent with this, an NLS prediction tool (29) identified this region as containing a putative NLS (data not shown). Since lysine- and arginine-rich motifs are also characteristic of NoLS, we further scanned the USP10 amino acid sequence using an NoLS prediction tool (30). This analysis revealed that the 139–166 region contains a putative NoLS that partially overlaps with the predicted NLS (Figure 7A). To test whether the 139–166 region functions as an NoLS, we fused amino acids 139–166 of USP10 to the C-terminus of mCherry (mCh_139–166) and examined its localization in transfected cells. The fusion protein, but not mCherry alone (mCh), colocalized with the nucleolus, as demonstrated by nucleolar staining with the RNA-binding dye Nucleolus Bright Green (Figure 7B). Furthermore, mCh_139–166 also colocalized with the nucleolar marker fibrillarin, supporting the conclusion that the 139–166 region functions as an NoLS (Figure 7C). To examine whether USP10 localizes to the nucleolus through the 139–166 region, we fused GFP to the N-terminus of FL USP10 and Δ139–166 USP10, stably expressed these constructs in USP10-KO HCT116 cells, and assessed their colocalization with the nucleolus (Figure 7D). Although USP10 was predominantly localized to the cytoplasm, a fraction of FL USP10 colocalized with the nucleolus, as revealed by nucleolar staining. In contrast, Δ139–166 USP10 showed no detectable nucleolar colocalization. Together with the finding that the 139–166 region is sufficient to direct mCherry to the nucleolus (Figure 7C), these results indicate that the 139–166 region is required and sufficient for the nucleolar localization of USP10.

**FIGURE 7.**
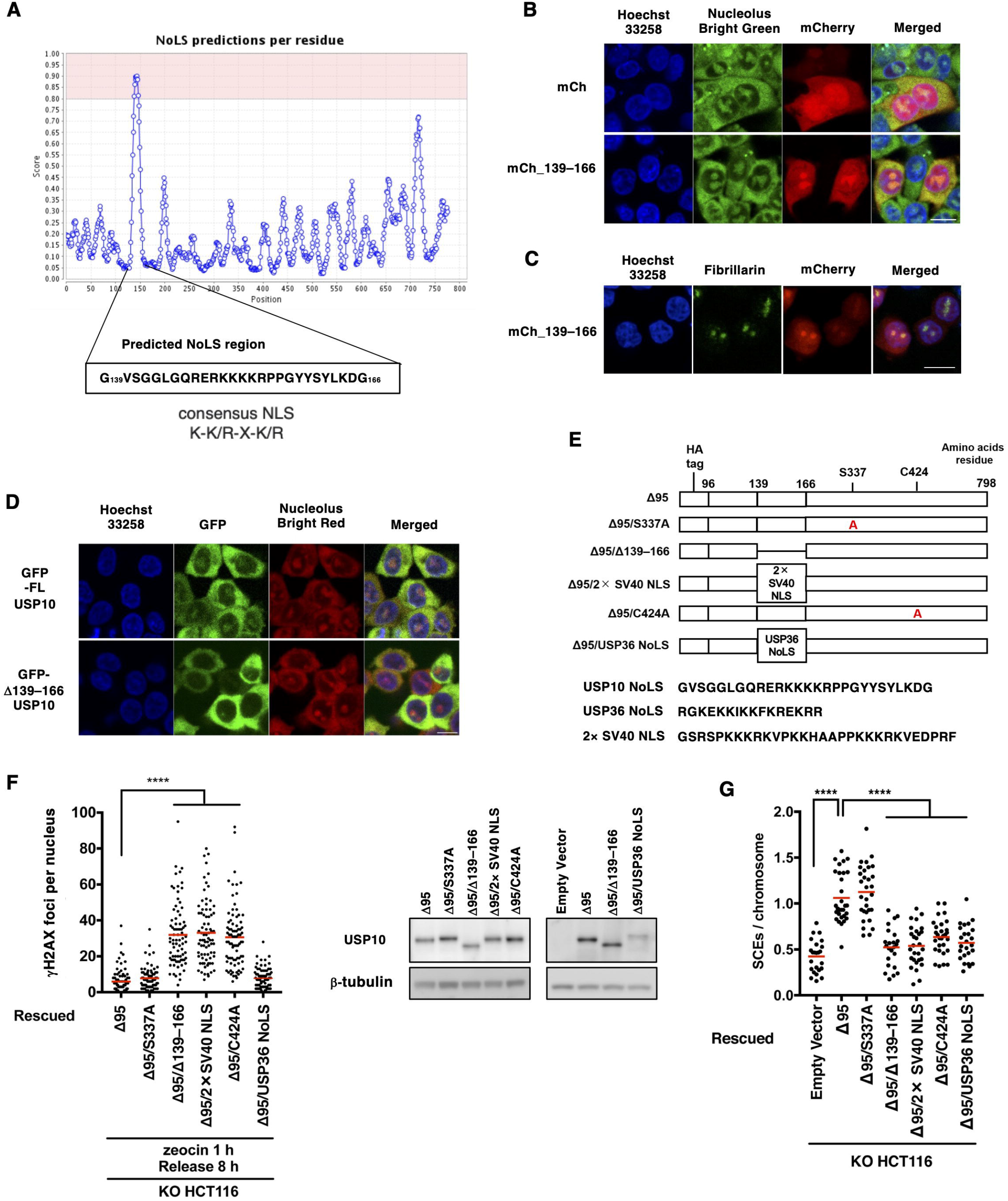
The USP10 139–166 region contains a nucleolar localization signal (NoLS). (A) Predicted NoLS sequence within USP10 and a consensus nuclear localization signal (NLS). (B) HCT116 cells were transfected with expression plasmids encoding mCherry (mCh) or mCherry fused to USP10 amino acids 139–166 (mCh_139–166) and stained with Nucleolus Bright Green. Nuclei were counterstained with Hoechst 33258. (C) HCT116 cells were transfected with the mCh_139–166 expression plasmid and stained with an anti-fibrillarin antibody. (D) USP10-KO HCT116 cells expressing GFP-fused FL USP10 or Δ139–166 USP10 were stained with Nucleolus Bright Red. Nuclei were counterstained with Hoechst 33258. (E,F) DSB repair was examined in USP10-KO HCT116 expressing the indicated USP10 mutants (E) following zeocin treatment. Expression of the mutants was determined by WB using an anti-USP10 antibody. Amino acid sequences of USP10 NoLS, USP36 NoLS, and 2× SV40 NLS are shown in (E). (G) SCE frequencies following CPT treatment in USP10-KO HCT116 expressing the indicated mutants. *P* values were determined by Dunnett’s multiple comparison test. ****, *P* < 0.0001.

We next assessed whether a heterologous NoLS could substitute for the USP10 NoLS. Specifically, we replaced the 139–166 region in the Δ95 USP10 mutant with the NoLS of USP36, a well-characterized nucleolar protein (31). If nucleolar localization alone is sufficient for USP10 function in DSB repair, this substitution would be expected to restore the repair capacity of the Δ139–166 mutant. In parallel, we tested whether the canonical SV40 NLS could functionally replace the USP10 NoLS. In addition, because ATM-mediated phosphorylation of serine 337 has been reported to promote USP10 nuclear translocation following DNA damage (10), we examined a phosphorylation-deficient mutant in which serine 337 was substituted with alanine (S337A) (Figure 7E). These constructs were introduced into USP10-KO HCT116 cells, and DSB repair was evaluated following zeocin treatment (Figure 7F). While the USP36 NoLS successfully restored DSB repair capacity in the Δ139–166 mutant, the SV40 NLS did not, indicating that nucleolar localization—but not nuclear localization per se— is required for USP10 to facilitate DSB repair. The S337A mutant retained DSB repair capacity comparable to the Δ95 mutant, suggesting that phosphorylation of serine 337 is not essential for this function. However, when SCE activity was assessed, the USP36 NoLS failed to substitute for the USP10 NoLS (Figure 7G), suggesting that the latter contributes to functions beyond nucleolar targeting, potentially related to HJ resolution leading to crossover.

## 4 Discussion

USP10 is a multifunctional deubiquitinase that regulates diverse cellular processes, including the DNA damage response. Here, we show that USP10 is required for HR-mediated repair of DSBs and for maintaining genomic stability. USP10-depleted cells exhibit spontaneous micronuclei, impaired DSB repair after zeocin and CPT treatment, and reduced SCE activity. They are also more sensitive to IR and MMC and display increased chromosomal abnormalities after MMC treatment. Re-expression of WT USP10—but not a catalytically inactive mutant—rescued these phenotypes, indicating that its deubiquitinase activity is required for HR-mediated DSB repair.

Mechanistically, our data place USP10 downstream of end resection and RAD51 filament formation. USP10-KO cells form RAD51 foci with normal kinetics after DSB induction, but these foci persist longer than in WT cells. This suggests that resection and RAD51 loading are intact, whereas a later step—such as strand invasion or RAD51 dissociation—is defective. Persistent RAD51 foci have been reported after depletion of several RAD51-regulatory factors (e.g., RFWD3, RAD54, FIRRM, and FIGNL1), which modulate RAD51 stability or promote filament disassembly (32-35). We therefore speculate that USP10 regulates HR by deubiquitinating one or more RAD51 regulators, thereby facilitating timely remodeling or removal of RAD51 nucleoprotein filaments to enable downstream recombination steps.

Recent studies have identified several DNA repair factors as USP10 substrates, including TRMT10A, which promotes BRCA1 recruitment during HR (36); XAB2, which facilitates ANXA2-mediated ICL repair (37); and PARP1, which facilitates HR repair (38). However, USP10 deletion did not reduce the steady-state levels of TRMT10A, XAB2, or PARP1 in HCT116 or MEF cells (Figure S7). These findings indicate that stabilization of these factors is not a general function of USP10 and does not explain the HR and ICL repair defects observed here. Instead, our data support a model in which USP10 regulates genome maintenance through additional, context-dependent mechanisms.

We identified a previously uncharacterized NoLS in USP10 (amino acids 139–166) that is required for HR repair. Replacing USP10’s NoLS with that from USP36 preserved DSB-repair capacity after zeocin treatment, supporting a role for nucleolar targeting in USP10 function. However, the USP36-NoLS swap did not fully restore SCE activity, indicating that nucleolar localization alone is insufficient for all USP10 functions in HR. The NoLS region may therefore have an additional role, for example in facilitating HJ resolution associated with crossover outcomes rather than HJ dissolution or SDSA.

How nucleolar localization enables USP10 to promote HR remains unclear. Recent work suggests that pre-rRNA can act as a scaffold for recruitment of RAD51AP1 to DSBs and facilitate HR (39). One possibility is that USP10 regulates nucleolar pathways involved in ribosome biogenesis—for example, by controlling ribosomal protein stability, rRNA transcription, or maturation—and that disruption of these processes in USP10-deficient cells perturbs pre-rRNA homeostasis and impairs HR. In this model, nucleolar proteins involved in ribosome biogenesis may represent additional USP10 substrates alongside RAD51 regulators.

HR defects predispose to cancer (40), and the genomic instability observed in USP10-deficient cells supports a tumor-suppressive role for USP10 in some contexts (41). Conversely, USP10 has also been reported to promote tumorigenesis and therapy resistance, in part through DNA repair pathways (41). Thus, USP10-dependent HR may contribute to context-dependent cancer phenotypes and represents a potential therapeutic target.

## Supporting information

Supplementary doc S1

## Abbreviations

HJ: Holliday junction
HR: homologous recombination
NHEJ: non-homologous end joining
NoLS: nucleolar localization signal
SCE: sister chromatid exchange
SDSA: synthesis-dependent strand annealing

## Acknowledgments

We thank Edanz (https://jp.edanz.com/ac) for editing a draft of this manuscript.

## Disclosure

### Funding Information

This work was supported by Grant for Promoted Research from Kanazawa Medical University (S2017-10) and JSPS KAKENHI Grant Numbers JP18K06070, JP22K12382.

### Conflict of Interest

The authors have no conflict of interest.

### Ethics Statement

- Approval of the research protocol by an Institutional Reviewer Board: N/A
- Informed Consent: N/A
- Registry and the Registration No. of the study/trial: N/A
- Animal Studies: N/A

### Author Contributions

**Koichi Utani:** data curation, investigation, writing–original draft. **Ryo Sakasai:** supervision, writing–review and editing. **Takako Okuwa:** supervision. **Toshiki Himeda:** supervision. **Kuniyoshi Iwabuchi:** supervision, resources. **Masaya Higuchi:** conceptualization, writing–review and editing.

## List of Supporting Information

Doc. S1

Figures S1–S7

## Notes

### Competing Interest Statement

The authors have declared no competing interest.

