## Supplementary doc S1 for "USP10 Facilitates Homologous Recombination-Mediated DNA Double-Strand Break Repair through Localization to the Nucleolus"

**Doc S1. Supplementary Materials and Methods**

**Cell Culture**

A mouse embryonic fibroblast (MEF) cell line harboring floxed *Usp10* exon 3 alleles (*Usp10* floxed MEFs) (1) was established using the conventional 3T3 method. MEFs and the human cell lines HCT116, MCF-7, WI-38, and 293T were cultured in DMEM supplemented with 10% FCS, 100 U/mL penicillin and 100 μg/mL streptomycin (FUJIFILM Wako Chemicals, Japan). For MEF cultures, 55 μM 2-mercaptoethanol (Thermo Fisher Scientific, USA) was added. All cell lines were maintained at 37 °C in a humidified atmosphere containing 5% CO2.

**Plasmids**

pCAG-nCreGFP, an expression vector encoding GFP-fused Cre recombinase, was a kindly provided by Dr. Hirohide Takebayashi (Kyoto University). The control vector pCAG-GFP was generated by inserting EGFP from pEGFP-N3 (Clontech Takara, Japan) into the EcoR I and Not I sites of the pCAG vector. Entry vectors for mouse USP10 and its mutants have been described previously. Human USP10 cDNA was PCR-amplified from a cDNA library generated from HeLa cells using the In-Fusion SMARTer directional cDNA library construction kit (Clontech Takara) and cloned into pENTR/D-TOPO (Thermo Fisher Scientific). Entry vectors encoding HA-tagged human USP10 mutants were generated by PCR or were kindly provided by Dr. Masahiko Takahashi (Niigata University). pcDNA3.1_AGIA-AirID was kindly provided by Dr. Tatsusya Sawasaki (Ehime University). The AGIA-AirID fragment was PCR-amplified and inserted into pENTR-Δ138 USP10 using the In-Fusion cloning method (Clontech Takara). Additional deletion mutants derived from AirID-Δ138 USP10 were generated by PCR. To construct an entry vector encoding mCherry fused to the USP10 nucleolar localization signal (NoLS), the mCherry fragment was first PCR-amplified from pmCherry-N1 (Clontech Takara) and cloned into pENTR/D-TOPO. The region corresponding to USP10 amino acids 139–166 region was then PCR-amplified and inserted into the Xho I and Spe I sites downstream of mCherry. Entry vectors encoding GFP-fused USP10 and USP10 Δ139-166 were generated by inserting a PCR-amplified GFP fragment into their respective entry vectors using the In-Fusion cloning method. Replacement of USP10 NoLS with the USP36 NoLS or 2× SV40 NLS was performed using the In-Fusion method, with annealed oligonucleotides for the USP36 NoLS or a PCR-amplified fragment from pCBASceI (Addgene 26477) for 2× SV40 NLS. A lentiviral destination vector, CSII-CMV-RfA-IRES-Bsd, was constructed by inserting the RfA cassette (Thermo Fisher Scientific) into the Xho I site of CSII-CMV-MCS-IRES-Bsd (RIKEN BRC, Japan). USP10 cDNA and its derivatives were transferred into CSII-CMV-RfA-IRES-Bsd using LR Clonase (Thermo Fisher Scientific). SGEP, a lentiviral vector for shRNA mediated knockdown (KD) (2), was kindly provided by Dr. Johannes Zuber (Research Institute of Molecular Pathology). The target sequences used were as follows: human USP10, 5’-TTGGAGTTAAAATGTTAGTCTA-3’; and *Renilla* luciferase control (Ren), 5’- TAGGAATTATAATGCTTATCTA -3’.

**Lentiviral Transduction**

For lentivirus production, 293T cells were transfected with the lentiviral vector together with packaging plasmids (pCAG-HIVgp and pCMV-VSV-G-RSV-Rev; RIKEN BRC) using PEI MAX (Polysciences, USA) and cultured for 72 h. The resulting culture supernatant was collected and added to target cells in the presence of 8 μg/mL polybrene (Sigma, USA). Transduced cells were subsequently selected with the appropriate antibiotics: 1 μg/mL puromycin (InvivoGen, USA) or 10 μg/mL blasticidin S (InvivoGen) for HCT116 cells, and 4 μg/mL blasticidin S for MEF cells.

**Establishment of USP10 KO or KD Cell Lines**

To establish USP10 KO and control WT MEFs, *Usp1*0 floxed MEFs were transfected with pCAG-nCreGFP or pCAG-GFP, respectively, using FuGENE 6 (Promega, USA). EGFP-positive cells were sorted using a FACSAria II cell sorter (Becton Dickinson, USA). HCT116 USP10 KO cell clones were generated by transfection with pGuide-it-ZsGreen1 (Clontech Takara) containing a gRNA sequence (5’-CTTACCTCAACTGAAGATCG-3’) using PEI MAX, followed by limiting dilution. For USP10 KD, cells were infected with lentiviruses produced from SGEP plasmids and selected with puromycin.

**DNA Damage Induction**

Cells were treated with zeocin (Thermo Fisher Scientific), etoposide (Sigma), camptothecin (CPT; Sigma), and mitomycin C (MMC; FUJIFILM Wako Chemicals). For UV-mediated DNA double-strand break (DSB) induction, cells were first labeled with 10 μM BrdU (Sigma) for 72 h and then irradiated with UV at a dose of 3 mJ/cm^2^ using a FUNA-UV-LINKER (Funakoshi, Japan). X-ray irradiation was performed using a M-80WE device (SOFTEX, Japan). SCR7 (Sigma) was used as an inhibitor of DNA ligase IV.

**Antibodies**

For western blotting, the following antibodies were used: USP10 (1:1000; CST 8501, USA or Proteintech 99374-1-AP, USA), VCP (1:3000; CST 2648), SAPS1 (1:1000; Bethyl A300-968A-T, USA), RPS3 (1:400; CST 2579), AGIA tag (1:10000; a kind gift from Dr. Sawasaki, Ehime University), G3BP1 (1:5000; Proteintech 13057-2-AP), TRMT10A (1:2000; Proteintech 17291-1-AP), XAB2 (1:4000; Proteintech 10637-1-AP), PARP1 (1:1000; CST 9542), HA (1:10000; Roche 12158167001, USA), anti-Rat IgG-HRP (1:10000; Santa Cruz sc-2006, USA), and HRP-conjugated anti-mouse and anti-rabbit IgG (1:10000; Bio-Rad 170-6516 and 170-6515, USA, respectively). For immunostaining, the following antibodies were used: γH2AX (1:1000; Millipore 05-636 or 07-164, USA), RAD51 (1:200; Santa Cruz sc-8349), RAD51 (1:1000; Abnova H00005888-B01P, Taiwan), Alexa Fluor 594-conjugated goat anti-mouse IgG, and Alexa Fluor 488-conjugated goat anti-rabbit IgG (1:1000; Thermo Fisher Scientific A11005 and A11008, respectively).

**Immunostaining**

Cells cultured on coverslips were fixed in PBS containing 4% paraformaldehyde (PFA) for 10 min, followed by permeabilization with methanol at −20 °C for 10 min and with 0.5% Triton-X-100 in PBS at 4 °C for 10 min. Cells were then incubated with the indicated primary antibodies, followed by incubation with fluorescence-labeled secondary antibodies. Nuclei were stained with Hoechst 33258 (Thermo Fisher Scientific) or DAPI (Sigma). Nucleoli were visualized using Nucleolus Bright Green or Red reagents (Dojindo, Japan). Images were acquired using an LSM710 confocal microscope (Zeiss, Germany) and analyzed using Fiji software.

**Western Blotting**

Total cell lysates were prepared by lysis in 1× SDS sample buffer. Western blot analysis was performed as previously described (3). Can Get Signal solution (TOYOBO, Japan) was used for antibody incubation

**Neutral Comet Assay**

A comet assay kit (Trevigen, USA) was used according to the manufacturer’s protocol. Slides were stained with SYBR Gold (Thermo Fisher Scientific). Images of stained nuclei were acquired using an IX73 microscope (Olympus, Japan), and more than 80 nuclei per sample were per sample were analyzed for olive tail moment using CometScore software (TriTeK, USA).

**MTS Cell Viability Assay**

For MTS assay, 2,500 cells were seeded in a 96 well plate, treated with the indicated drugs, and cultured for 36 h. Cells were then incubated with CellTiter 96 AQueous One Solution Cell Proliferation Assay reagent (Promega) for 1 h. Cell viability was accessed by measuring absorbance at 490 nm using a Multiskan GO microplate reader (Thermo Fisher Scientific).

**Clonogenic Colony Formation Assay**

MEF cells (500 cells) were plated in 6-cm dishes or 6-well plates and then exposed to X-ray irradiation or treated with MMC, respectively. After 8–11 days, colonies were fixed with 2% PFA in PBS and stained with crystal violet.

**Sister Chromatid Exchange (SCE) Assay**

The SCE assay was performed as previously described (4, 5). Briefly, cells were labeled with 10 μM BrdU for two cell-cycle doubling times, as illustrated in Fig. 3E, and treated with 10 nM CPT during the final 8 h of labeling to induce DSBs. To enrich for mitotic cells, 10 μg/mL colcemide (Thermo Fisher Scientific) was added for 1.5 h prior to fixation. Cells were then fixed with Carnoy’s fixative (methanol:acetic acid, 3:1), and metaphase chromosome spreads were prepared. Slides were stained with 2 μg/mL Hoechst 33258 in 2× SSC, rinsed sequentially with 2× SSC and 0.5× SSC, and exposed to sunlight for 2–4 h. Slides were subsequently incubated in 0.5× SSC at 58 °C for 10 min, washed with distilled water, and stained with 5% Giemsa in 0.05× SSC for 1 h. Slides were then briefly destained in 0.5× SSC and rinsed with distilled water. For SCE quantification, images of 30–50 metaphase spreads were acquired using an Eclipse E600 microscope (Nikon, Japan), and SCE events were counted using Fiji software.

**Analysis of Chromosomal Aberrations**

Cells were treated with 0 or 50 nM MMC for 3 days. Metaphase chromosome spreads were prepared as described above for the SCE assay. After complete air-drying, slides were washed in 2× SSC, desalted, and stained with Giemsa solution. Fanconi anemia (FA)-specific chromosomal aberrations were evaluated according to previously defined criteria (6). Briefly, single gaps and breaks were each counted as one event, whereas triradial and quadriradial structures were each counted as two events. FA-type chromosomal aberrations were scored per metaphase, with approximately 30 metaphase spreads analyzed for each cell line and treatment condition.

**Statistical analysis**

GraphPad Prism software, version 7 (GraphPad Software), was used for all statistical analyses.
