## Supplementary figures for "USP10 Facilitates Homologous Recombination-Mediated DNA Double-Strand Break Repair through Localization to the Nucleolus"

**FIGURE S1.** Generation of USP10-KO MEFs. **(A)** Genomic structure of mouse *Usp10*. Black triangles indicate loxP sites flanking exon 3. The conditional *Usp10*$^{flox}$ allele was converted to the recombined *Usp10*$^{\Delta}$ allele by Cre-mediated recombination. Positions of primers used for genotyping are indicated by small arrows. **(B)** PCR genotyping of USP10-KO MEFs. Primer sequences were 5’-GGTGTTTGGGGCTCGGTTCTGTCA-3’ and 5’-TGGCAGTTGTGGTGGTTTGAGTATG-3’. **(C)** USP10 protein expression in WT and USP10-KO MEFs. Total cell lysates were prepared from WT and USP10-KO MEFs and analyzed by western blotting (WB) using anti-USP10 or anti-TSC1 antibodies. TSC1 was used as a loading control.

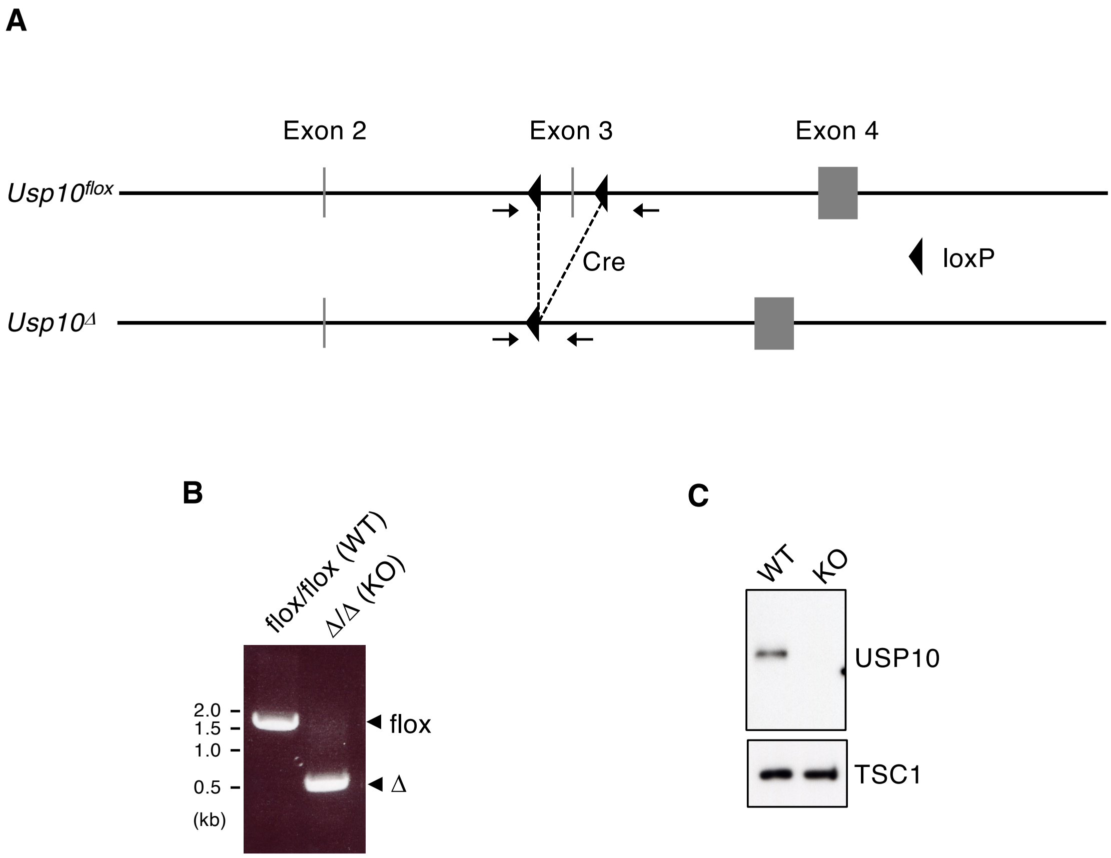

**FIGURE S2.** USP10 is required for DSB repair in human cell lines. **(A)** USP10 knockdown (KD) in HCT116, MCF-7, and WI-38 cells. shRNA targeting *Renilla* luciferase (Ren) was used as a control. USP10 expression was determined by WB using an anti-USP10 antibody. β-tubulin was used as a loading control. **(B–D)** γH2AX foci formation in control and USP10-KD human cell lines following zeocin treatment. **(E,F)** USP10 expression (E) and γH2AX foci formation (F) in USP10 KD-HCT116 cells reconstituted with WT USP10 or the C428A mutant. **(G)** USP10 expression in USP10-KO HCT116 cell clones compared with that in WT and USP10-KD cells. VCP was used as a loading control. **(H,I)** Micronuclei formation (H) and γH2AX foci formation following zeocin treatment (I) in USP10-KO HCT116 cells. *P* values were determined by the Mann–Whitney *U* test (B–D) and Dunnett's multiple comparison test (F,H). ns, not significant; **, $P < 0.01$; ****, $P < 0.0001$.

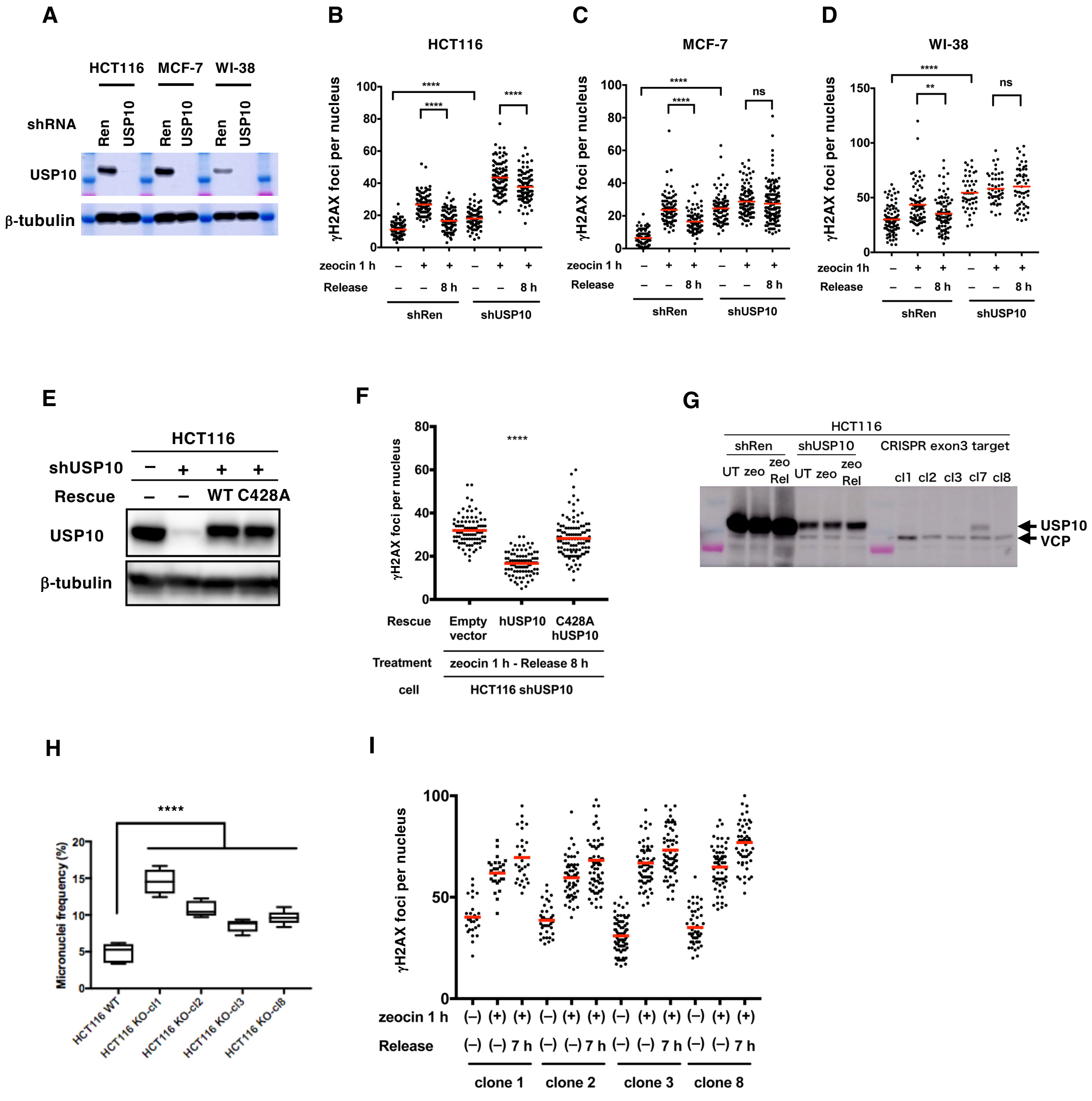

**FIGURE S3.** USP10-KO MEFs exhibit reduced colony formation following DNA damage. **(A,B)** Clonogenic colony formation assays of WT and USP10-KO MEFs following X-ray irradiation (A) or MMC treatment (B). *P* values were determined by Student's *t* test. ***, $P < 0.001$; ****, $P < 0.0001$.

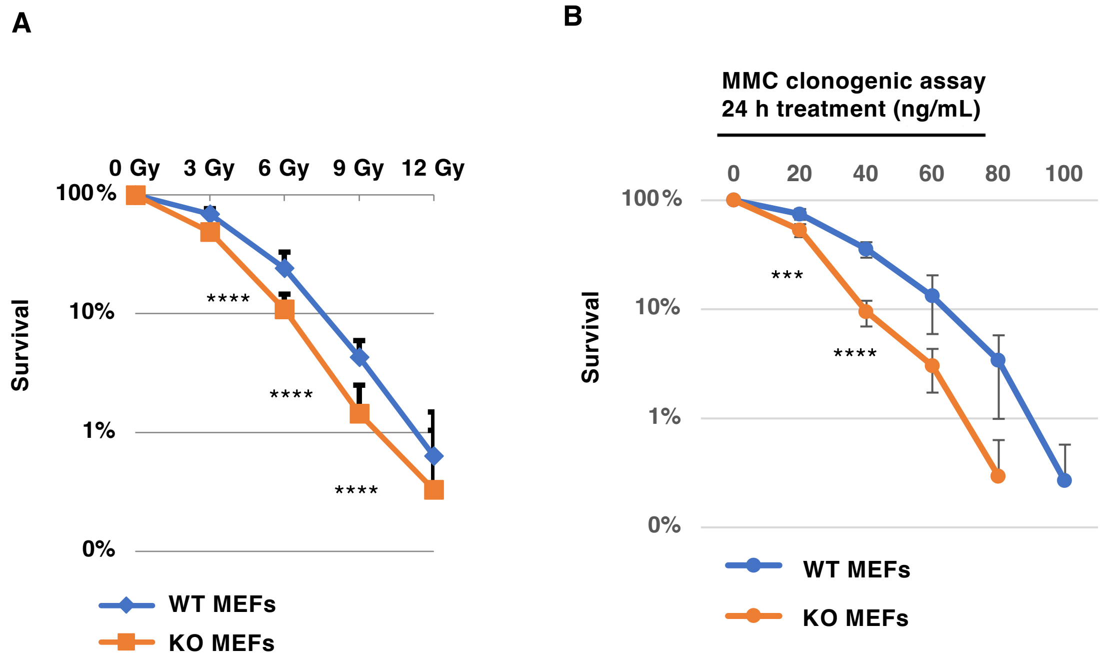

**FIGURE S4.** Persistent RAD51 foci formation in USP10-KO HCT116 cells. **(A)** RAD51 (green) and γH2AX (red) foci formation in WT and USP10-KO HCT116 cells following treatment with 100 μg/mL zeocin. Nuclei were counterstained with DAPI. Scale bar, 10 μm. **(B)** Quantification of RAD51 foci. *P* values were determined by the Mann–Whitney *U* test. ****, $P < 0.0001$.

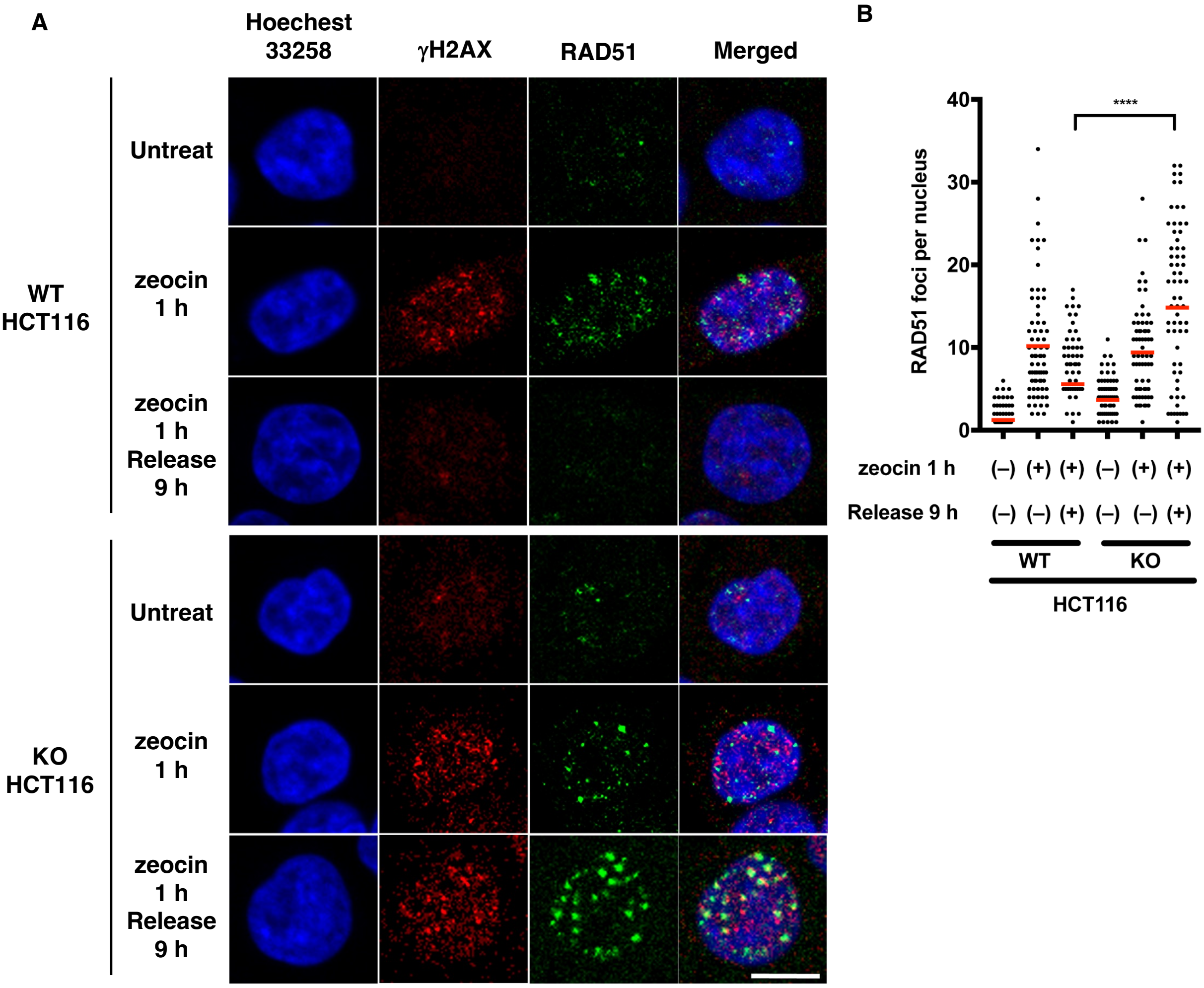

**FIGURE S5.** Localization of USP10 and γH2AX in USP10-KO HCT116 cells. USP10 (green) and γH2AX (red) localization in USP10-KO HCT116 cells expressing WT USP10 or the C424A mutant following UV irradiation, with or without BrdU labeling. Nuclei were counterstained with Hoechst 33258. Scale bar, 50 μm.

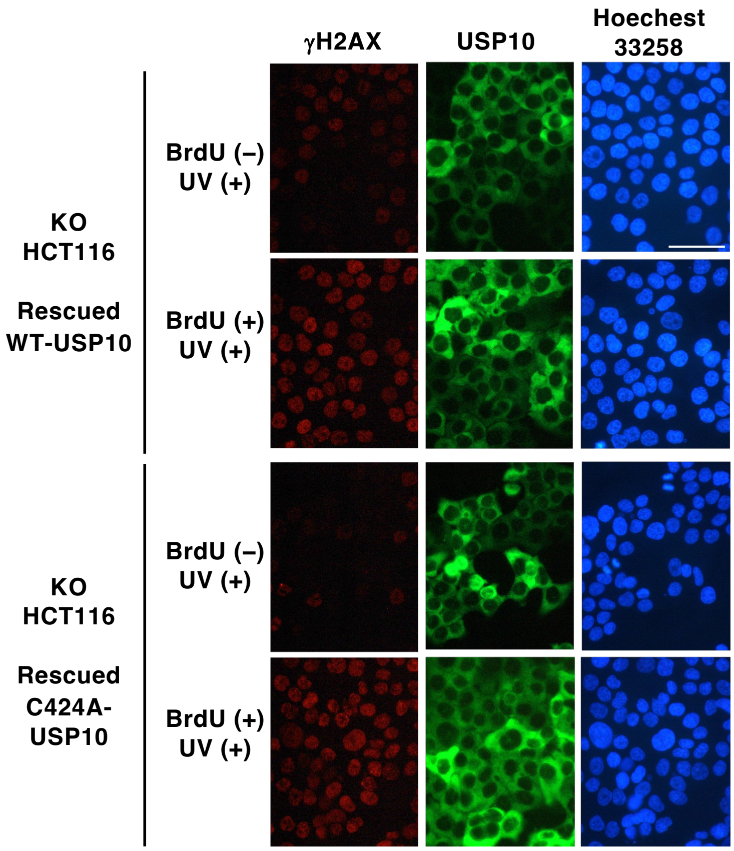

**FIGURE S6.** The USP10 region 166–383 is not required for DSB repair. **(A)** USP10-KO HCT116 cells were transduced with AirID or AirID-fused USP10 Δ138 or Δ166 mutants, and DSB repair following zeocin treatment was examined. Expression of the mutants was determined by WB using an anti-AGIA tag antibody. G3BP1 was used as a loading control. **(B)** USP10-KO HCT116 cells were transduced with the indicated AirID-fused USP10 mutants, and DSB repair following zeocin treatment was examined. Expression of the mutants was determined by WB using an anti-AGIA tag (upper right) or anti-USP10 antibody (lower right). SAPS1 or β-tubulin was used as a loading control. *P* values were determined by Dunnett's multiple comparison test. ns, not significant; ****, $P < 0.0001$.

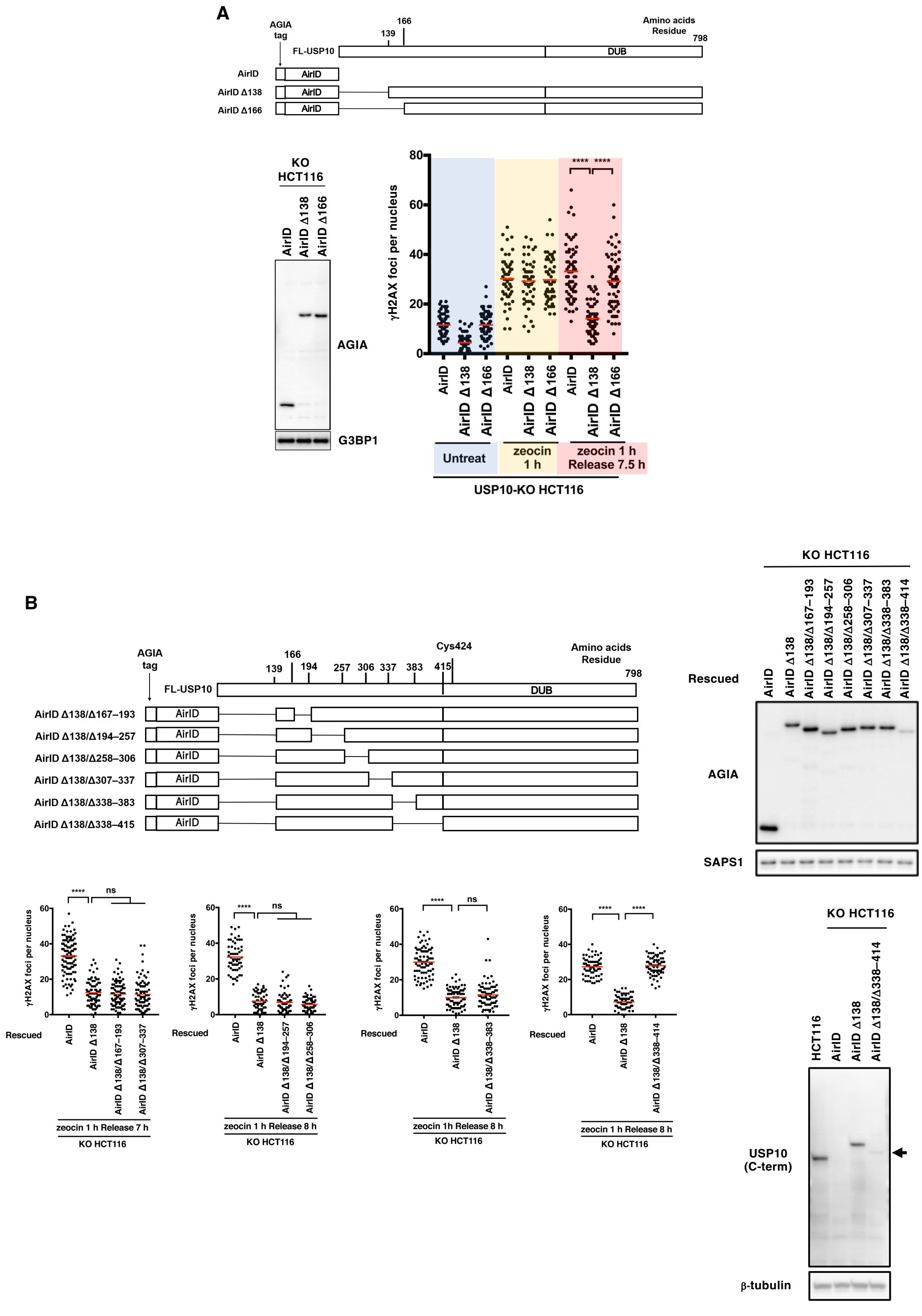

**FIGURE S7.** The protein levels of TRMT10A, XAB2, and PARP1 are unchanged in USP10-KO HCT116 cells and USP10-KO MEFs, as determined by WB using anti-TRMT10A, anti-XAB2, and anti-PARP1 antibodies. β-tubulin was used as a loading control.

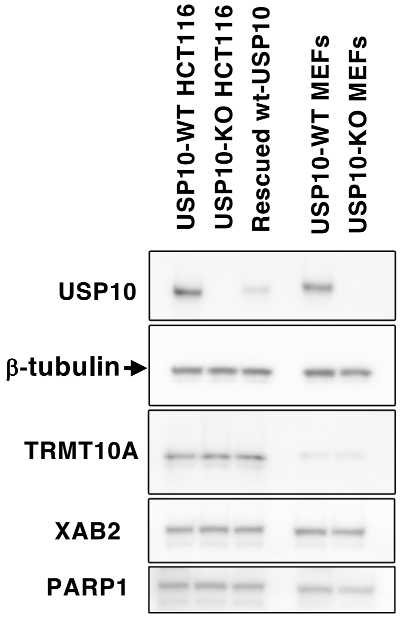
